# Annotation of 200 Insect Genomes with BRAKER for Consistent Comparisons across Species

**DOI:** 10.1101/2025.04.17.649312

**Authors:** Stepan Saenko, Katharina J. Hoff, Mario Stanke

## Abstract

The annotation of genomes lags behind their sequencing and assembly. For example, of 5,092 insect species in GenBank, as one of the most widely used databases, only 375 currently have annotated genomes within this database.

Additionally, species that were previously annotated can benefit from reannotation using RNA-Seq and protein data that have been added since their last annotation, as well as from state-of-the-art annotation methods, whose accuracy has improved. Heterogeneous annotations performed with different tools and protein databases can introduce artifactual differences when comparing gene sets or gene structures between species.

Recently, the BRAKER3 annotation pipeline was introduced that integrates evidence from RNA-Seq and from a protein database. It was benchmarked as one of the most accurate annotation methods. Here, we introduce an automated genome annotation workflow that allows to annotate a list of species with BRAKER3 and VARUS for RNA-Seq retrieval, or in the absence of transcriptome data, with BRAKER2, with minimal manual intervention. We selected a diverse set of 200 insect species from different families, including 85 species previously lacking annotations in GenBank. Using currently available RNA-Seq and protein sequence data, we applied our automated workflow to annotate these genomes and conducted downstream analyses typically performed in comparative genomics studies.

We present the resulting gene structures, protein sequences, gene ontology terms, orthologous gene groups and a species tree.

## Background & Summary

Over the past two decades, the number of sequenced insect genomes has increased dramatically, from just twenty species twenty years ago to over 6,000 today, according to GenBank statistics [1]. Additionally, other large-scale initiatives and external databases, such as Tree of Life [2], contribute to this growing pool of genomic data. However, having access to raw data is only the first step—accurate and comprehensive genome annotations are essential to address key biological questions and challenges in phylogenetics, comparative genomics, evolutionary biology, the analysis of gene functions and the study of developmental pathways.

While the number of insect species with scaffold- or chromosome-level assemblies in GenBank has risen to 3,062 as of January 2025, the percentage of species with annotations at GenBank has decreased to only 10%, underscoring a significant annotation bottleneck (Figure 1).

**Figure 1:**
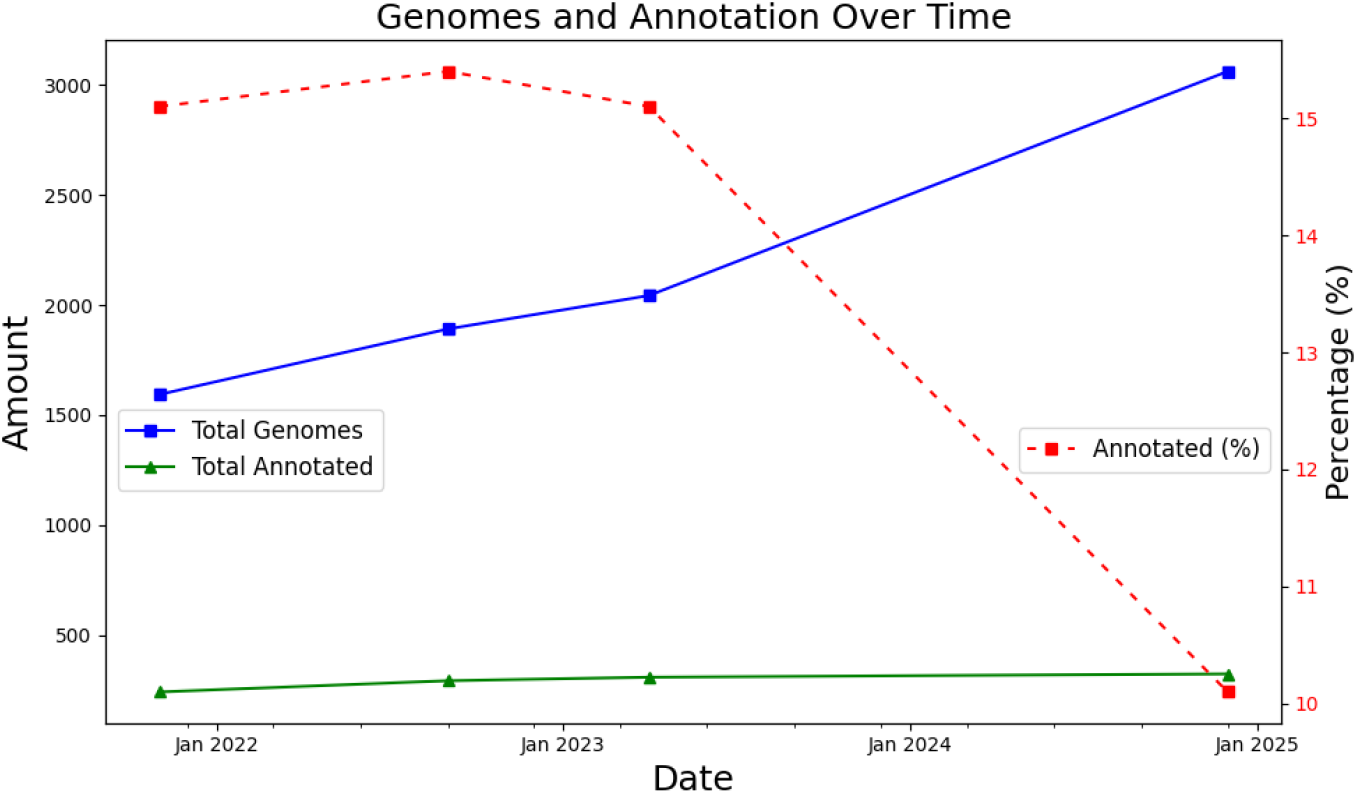
Total number of insect species in GenBank with at least scaffold level assembly and the percentage that has an annotation at GenBank

To maximize the benefit of genomic data, annotation pipelines must meet several criteria: algorithms must be up-to-date, raw data should be accessible, and the entire process must be reproducible; additionally, annotations must be high quality, standardized, and comparable between species. Comparability, in this context, means that observed genomic differences, such as gene truncations, the presence or absence of exons or even whole genes, or variations in gene structure, should reflect true biological distinctions rather than errors or missing genes or splice forms introduced by differing annotation methods or the use of other data sets (e.g., proteins or RNA-Seq).

Algorithm development has recently been shown to lead to improved accuracy [3–5], in addition to the progress that can be achieved when applying old algorithms to additional data. For example, the average increases in F1 scores when identifying complete protein-coding transcripts in eleven species were 4 percent points from BRAKER1 to BRAKER2 and an additional 24 percent points from BRAKER2 to BRAKER3. The latter achieved an average transcript-level F1 score of about 60%. Unfortunately, the current state of eukaryotic genome annotation is still not precise enough to ensure that all, or even most, of the structural differences identified between the annotations of closely related species can be reliably interpreted as biological variations. In addition, the problem of false positive differences can be exacerbated when annotations are generated by a variety of methods. For example, such heterogeneity in annotation methods has been shown to dramatically increase the apparent number of lineage-specific genes [6].

The recently developed BRAKER3 [4] integrates evidence from RNA-Seq in addition to evidence from a protein database, which was already integrated by BRAKER2 [7]. In both these run modes, the BRAKER pipeline automatically trains the parameters of hidden Markov models using the genome and the provided evidence and predicts protein-coding genes using the evidence again, including alternative splice forms. BRAKER3 was benchmarked under controlled conditions on the similarity between database and target proteins on 11 species, including *Drosophila melanogaster, Bombus terrestris* and *Parasteatoda tepidariorum*. It performed best among the pipelines compared [4].

The BRAKER pipeline can be started with a single command line and, contrary to MAKER [8], for example, does not require manual steps. However, the RNA-Seq data and a protein database have to be provided by a user and the input genome should be repeat masked. In addition, it is advisable to perform quality control steps after annotation, such as the execution of BUSCO [9] and OMArk [10]. As such preprocessing and postprocessing steps to annotation can require substantial manual effort if done for a large number of species, we have developed an automated *VARUS-BRAKER workflow* that performs them automatically and integrates multiple tools: VARUS [11], which automatically collects, selects, and aligns RNA-Seq data; repeat masking; BRAKER2 [7] and BRAKER3 [4], as well as downstream tools for quality control. As a result, the new VARUS-BRAKER workflow used in this study requires minimal manual effort per genome. In its user-friendliest run mode, only the binomial names of the species need to be input.

For the insect annotation data presented here, we focused on Holometabola and a diverse set of their outgroups, as our goal was the study of evolutionary innovations along the branch leading to the most recent ancestor of Holometabola. However, the annotation data and the orthologous groups of genes presented here can serve as a foundation for diverse other studies of insect genomics.

Holometabolous insects exhibit a unique larval stage that is morphologically distinct from the adult form. They undergo a complex transformation, with extensive reorganization and dedifferentiation of larval organs during the prepupal and pupal stages [12]. In contrast, hemimetabolous insects undergo a more gradual developmental process: their embryogenesis produces an adult-like larva that transitions into the adult form through a series of molts, with wings and genitalia typically appearing in the final molt. The origin of complete metamorphosis dates back approximately 400 million years [13] to the early Devonian period, an era marked by significant evolutionary innovations, including the emergence of winged insects (Pterygota), the advent of insect flight, and the development of holometaboly – an adaptation that likely contributed to the evolutionary success of these insects.

We here present whole-genome annotations of the protein-coding genes of 200 insect species, 85 of which did not have annotations in GenBank. Gene structures were predicted with BRAKER. RNA-Seq data was integrated for all species where it was available as well as evidence from protein homology. All 4,259,838 identified proteins were functionally annotated with GO terms. We identified groups of orthologs in the whole set of 200 proteomes, and subsequently constructed multiple sequence alignments of these protein families and a maximum likelihood species tree.

## Methods

The primary goal of this study was to address the gap in insect genome annotations by employing state-of-the-art tools, ensuring consistent annotation quality. Using the BRAKER3 pipeline [4], our objective was to minimize artifacts arising from heterogeneous annotation methods and incorporate the most current data, including protein and RNA-Seq datasets. To highlight the need for updates, we assessed the age of existing annotations using metadata from the NCBI RefSeq database, which provides reliable timestamps. Our analysis revealed a wide range of submission dates, with many annotations dating back several years. Specifically, out of the dataset examined, approximately 75% were submitted prior to 2022. The genome annotations presented in this work were generated using publicly available genome, RNA-Seq, and protein datasets. The analysis was carried out in three key stages: input specification, structural genome annotation, and downstream processing. An overview of the analysis workflow is shown in Figure 2.

**Figure 2:**
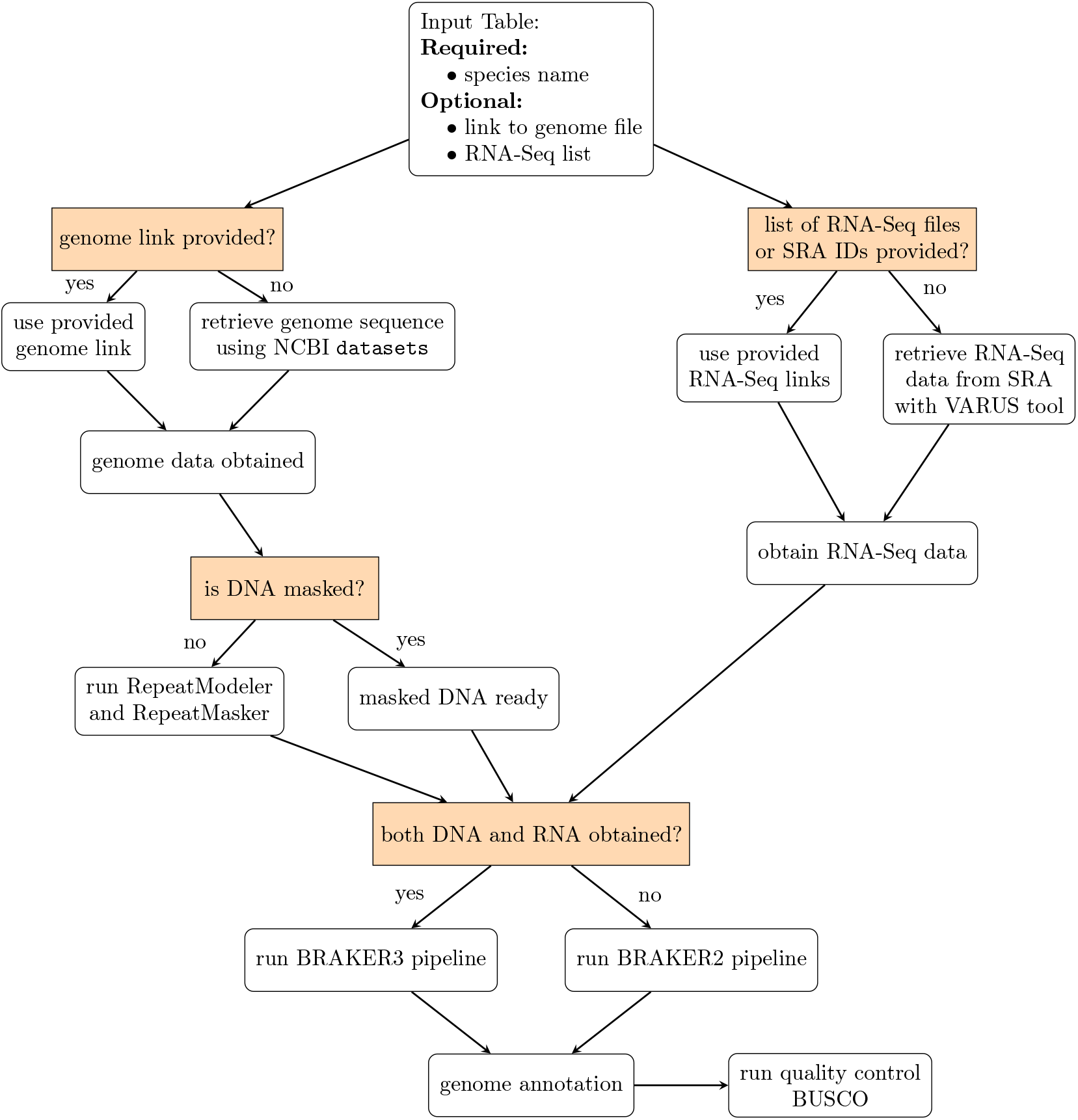
The scheme of the VARUS-BRAKER workflow.

In the first stage, *input specification*, the user prepares a text table with pointers to the genomes and optionally to the RNA-Seq data. Each line in the table specifies one genome to be annotated. There are several ways to specify the input data. As a minimum, when genome and RNA-Seq data are accessible from Genbank and Sequence Read Archive (SRA, https://www.ncbi.nlm.nih.gov/sra/), it is sufficient if a row just contains the species name, e.g. *Apis mellifera*. In general, the table has the following columns: species name (mandatory), optional link to the genome file (file path or URL) and optional list of links for RNA-Seq data or a list of SRA IDs. If there are no genome data provided, `datasets` from NCBI (v14.16.0) [14] will be used to download the assembly from GenBank, giving preference to assemblies tagged ‘reference’.

The second stage, *structural genome annotation*, encompasses all automatic steps up to and including the gene prediction. Initially, RepeatModeler2 [15] and RepeatMasker [16] generate a species-specific repeat library and soft-mask the genome. If no specific RNA-Seq data was specified, VARUS (v1.0.0) is used to automatically obtain RNA-Seq evidence. Varus uses NCBI’s `fastq-dump` to retrieve reads from SRA. The RNA-Seq reads are then aligned to the genome with HISAT2 [17]. The resulting SAM files are converted to BAM format, merged if multiple libraries were used, sorted, and indexed using SAMtools [18].

To obtain data to exploit homology with known proteins, the workflow automatically determines a suitable section of OrthoDB v12 [19] from the species name and using NCBI’s Taxonomy (https://www.ncbi.nlm.nih.gov/taxonomy/). This protein database is downloaded, unless already present, from https://bioinf.uni-greifswald.de/bioinf/partitioned_odb12. Then the gene models are called by the BRAKER pipeline (v3.0.3), which itself entails steps to train GeneMark-ETP [20] and AUGUSTUS [21] and predict evidence-based gene structures with them. If RNA-Seq data is available the BRAKER3 mode is used, otherwise the BRAKER2 mode to predict protein-coding gene structures from the soft-masked genome. The BRAKER3 mode requires RNA-Seq and is preferred as it was benchmarked to be significantly more accurate than BRAKER2 [4]. In BRAKER3 mode it utilizes GeneMark-ETP [22], which processes RNA-Seq alignments through StringTie2 [23] to assemble transcripts. GeneMarkS-T screens the assembled transcripts for potential genes. Additionally, DIAMOND [24] and GeneMark-EP+’s protein evidence pipeline are used to filter genes. GeneMark-ETP performs *ab initio* gene predictions based on self-training. Finally, AUGUSTUS is being trained on a reliable subset of predicted genes, and the final gene set is being consolidated using TSEBRA [25]. Protein evidence was incorporated during annotation, regardless whether BRAKER3 mode or BRAKER2 mode was used. This mode involves GeneMark-EP+, which self-trained GeneMark-ES to identify seed gene sequences. These sequences are then aligned to the protein database using DIAMOND, followed by accurate spliced alignment using Spaln2 [26]. The intermediate gene set generated by GeneMark-EP+ based on protein evidence was refined with AUGUSTUS.

Benchmark results comparisons to BRAKER3, when run with manually selected whole RNA-Seq libraries, are available in Section Technical Validation. Based on the experimental results, we chose a VARUS batch size of 75,000 bp and a maximum batch length of 600 bp. Increasing these parameters did not improve accuracy or precision; it only increased processing time. The optimal VARUS runtime was less than four hours.

In the third and final stage, *downstream processing*, the workflow performs quality assessment and functional annotation. BUSCO [9] is used to assess in particular the completeness of the predicted proteomes with regards to universal single-copy genes. In addition, FANTASIA [27] is used to determine functional annotations, integrating gene ontology terms and protein-domain annotations to enhance accuracy and completeness.

### Species Selection

As basis for selecting a sample of species, we used a phylogenetic tree derived from NCBI’s Taxonomy, afterwards we used the NCBITaxa module from the ete3 package [28] to handle taxonomy data, which included all insect species from the table at https://ftp.ncbi.nlm.nih.gov/genomes/GENOME_REPORTS/eukaryotes.txt with an assembly level higher than “contig,” resulting in a set of 3,062 species.

First, a subset of species was manually selected. The remaining species were then automatically selected to maximize diversity. Manual selection used two criteria: the number of citations in Google Scholar, reflecting their scientific relevance, and the availability of RNA-Seq data in the Sequence Read Archive (SRA), ensuring sufficient transcriptomic evidence for robust annotation. To obtain the remaining species, we developed and run a custom script that identifies for a given tree, set of manually selected leaves *A* and a given total number *n* a set *S* of leaves such that *A* and *S* together contain *n* leaves, and the tree restricted to *S* ∪ *A* and its ancestors has maximum total branch length, where the length of a branch is the number of taxonomic levels it spans. This script https://github.com/Gaius-Augustus/varus-braker/blob/main/scripts/maxSubtreeSet.pl is included in the BRAKER-VARUS repository on GitHub. This approach maximized diversity while including species that are suitable for comparative benchmarking or that are otherwise important. The choice also strikes a balance between reannotations (115) and annotations of previously unannotated genomes (85). Species with available RNA-Seq (169) were preferred over species without (31), as the accuracy of BRAKER is much higher in the former than in the latter [4].

With the above approach, initially, a total of 220 insect genome assemblies were retrieved from NCBI GenBank using the NCBI `datasets` [14] tool. The data set includes genomic data from 77 families across 14 orders. Later, we had to exclude some species from the data set due to their unsatisfactory predicted protein completeness level. All the remaining 200 species obtain 85% completeness. Nevertheless, we were able to maintain a balance between diversity and redundancy. Specifically, our taxon sampling includes the following species distributions: 40 from Lepidoptera, 34 from Hymenoptera, 24 from Hemiptera, and 20 from Coleoptera. A comprehensive list containing species, genus and accession numbers for the species’ genome assemblies, is provided in Supplementary Material 1.

### Structural genome annotation

All genome assemblies were processed with the VARUS-BRAKER workflow described above. In this use case with insects genomes, the workflow automatically retrieved the Arthropoda partition from OrthoDB v11 [19] from https://bioinf.uni-greifswald.de/bioinf/partitioned_odb11/Arthropoda.fa.gz.

The average running time of the workflow per species was approximately 20 hours, and 16 minutes, utilizing 36 CPUs, the runtime ranges from 5 h 09 min for *Goniozus legneri* to 78h 34 min for *Periplaneta americana*. This includes the time for automatic download of RNA-Seq data from NCBI, which varies due to the responsiveness of the SRA server.

### Orthology Analysis

The proteins predicted by BRAKER as part of the above workflow were grouped into orthogroups, representing sets of genes descended from a common ancestor. After annotating the 200 species listed in the Supplementary Table S1, we proceeded to find orthologous gene groups (orthogroups) in the whole set of 4,259,838 transcripts with OrthoFinder2 (v2.5.5) [29], which performed an all-versus-all comparison of protein sequences. OrthoFinder2 was executed with the following command:

~~~
python orthofinder.py -f species_proteins_dir/ -M msa -A mafft -t 36 -a 36
~~~

Here, the `-M msa` flag specifies that multiple sequence alignment (MSA)-based orthogroup inference is used, and `-A mafft` indicates that MAFFT is the chosen alignment tool. The options `-t 36` and `-a 36` specify the number of threads allocated for the analysis.

Along the orthogroup analysis protein multiple sequence alignments with MAFFT v7.505 [30] were constructed, the corresponding protein multiple sequence alignments files were used to construct a phylogenetic tree with RAxML Next Generation (RAxML-NG) [31], using the LG+G8+F substitution model. This model consists of the fixed LG empirical amino acid substitution matrix, empirical amino acid frequencies inferred from the alignment (+F), and rate heterogeneity modeled with 8 discrete categories of a gamma distribution (+G8). Phylogenetic inference was performed with 200 bootstrap replicates. The resulting tree (`200_insects_raxml.{pdf,nwk}`) and the input alignment (`SpeciesTreeAlignment.fa`) are provided in the data deposited at figshare https://figshare.com/s/fca8b91b7c3e1871d2ae.

The tree was built with the following command:

~~~
raxml-ng --all --model LG+G8+F/WAG+G8+F --tree pars{10} \
--bs-trees 200 --threads 36 \
--msa OrthoFinder2/Results/MultipleSequenceAlignments/SpeciesTreeAlignment.fa
~~~

Among the analyzed groups, all Endopterygota species were united into a hierarchical orthogroup, highlighting their shared evolutionary history.

The entire data processing workflow, including genome annotation, functional annotation, and orthology analysis, was implemented using standardized tools and versions, as documented in the Methods section. All software and parameters used for processing are provided in Supplementary Materials.

### Functional Annotation

Functional annotation was performed for all 200 species using the FANTASIA pipeline [27], which assigns Gene Ontology (GO) [32] IDs to transcripts. Multiple GO terms were assigned for some transcripts, reflecting their diverse functions. At the core of the FANTASIA pipeline is goPredSim[33], a similarity-based method for GO prediction.

FANTASIA provides Gene Ontology terms for each protein sequence. However, these identifiers are not human-readable. To enhance interpretability, we applied an R-based solution with a package `topGO` [34] for translating GO IDs into human-readable terms. GO terms are classified into three categories: Biological Process (BP), Molecular Function (MF), and Cellular Component (CC).

An example of applying this approach and analyzing the functional annotation data using GO terms is demonstrated in Figure 4. We obtained human-readable terms for the GO IDs and filtered by labels having insect, larva, or pupa in the name.

### Data Records

The dataset contains structural and functional annotations for 200 insect species. A taxonomic tree illustrating the relationships among these species is shown in Figure 3. The Orthofinder2 pipeline also constructs a species tree with the STAG method (Species Tree inferred from All Orthogroups). The tree topologies of this tree and the tree constructed using RAxML-NG have a Robinson-Foulds distance [35]) of 4.9%, i.e. 95.1% of of all possible splits that are induced by the edges of either tree are shared by the other tree. Both trees are available in the deposited data.

**Figure 3:**
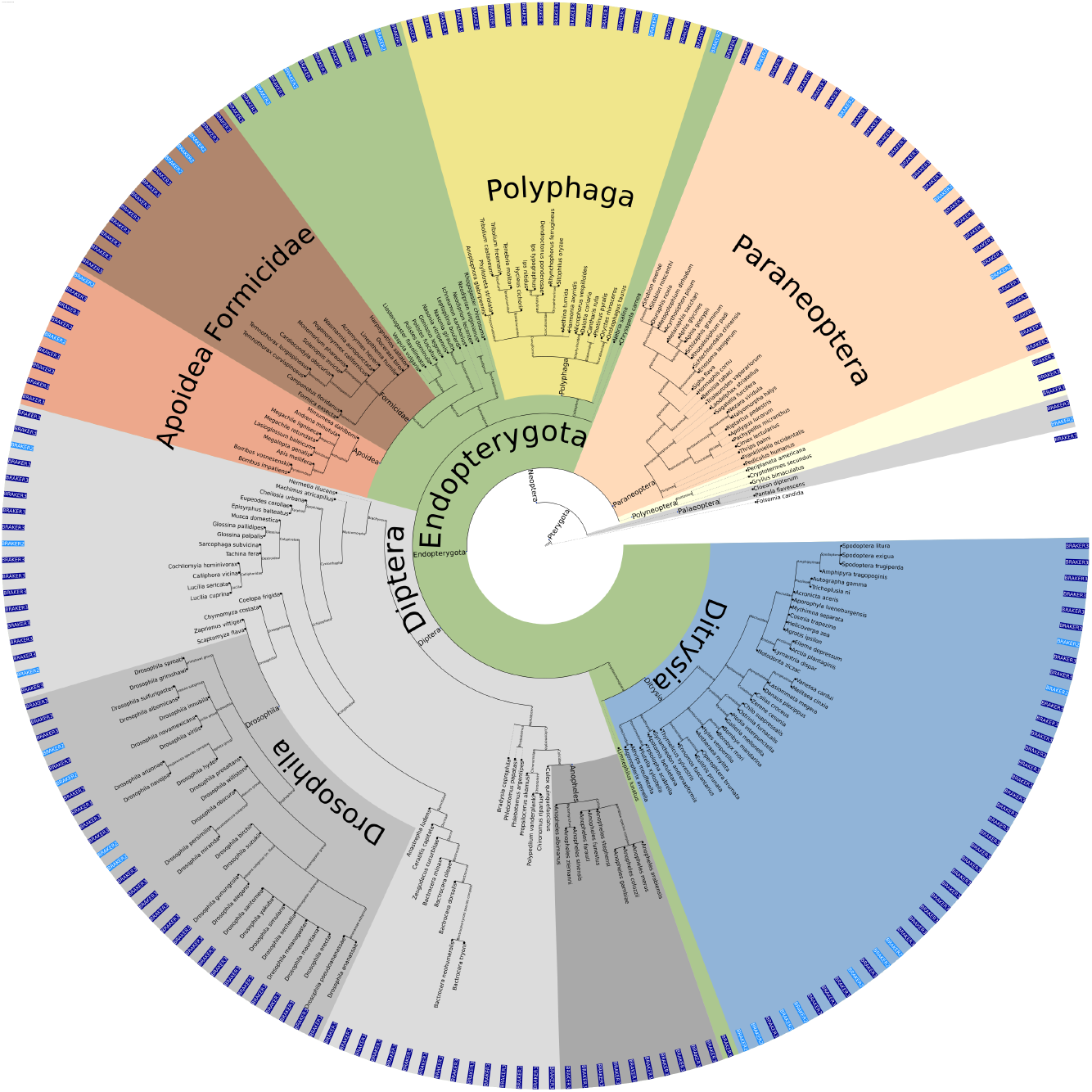
The taxonomy of the 200 selected species. A bright/dark blue label at the periphery of the circle marks whether annotations were done with BRAKER2 or BRAKER3, respectively.

**Figure 4:**
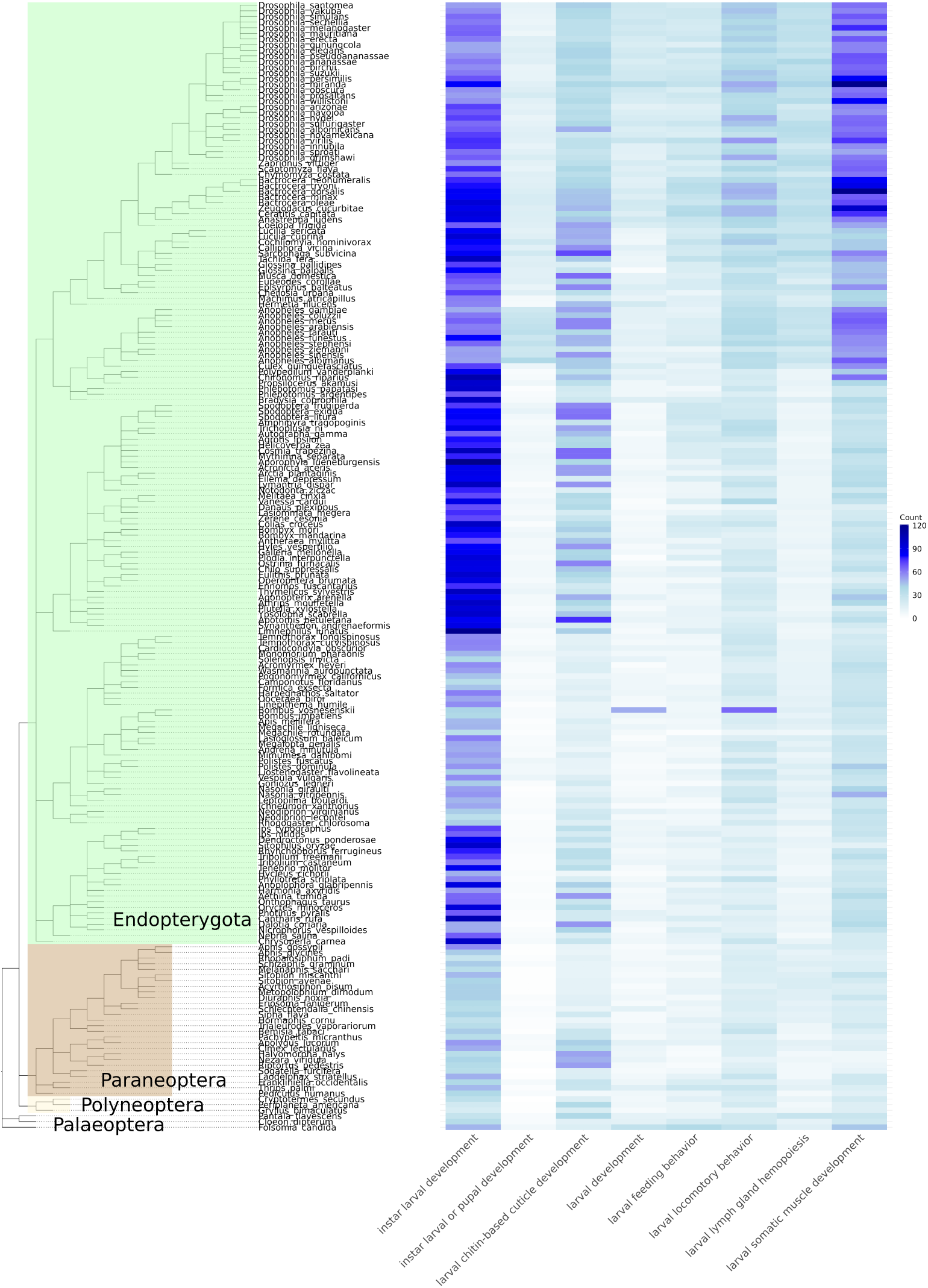
Visualization of an excerpt of the functional annotation. For selected biological processes as specified in gene ontology, the number of genes with this GO term annotation is shown.

Figures 5 and 6 show the distributions of genome sizes and protein lengths. The average genome size is 413,245,217 bp, and the sizes range between a minimum of 86 Mb *Propsilocerus akamusi* and a maximum of 3 Gb for *Periplaneta americana*.

**Figure 5:**
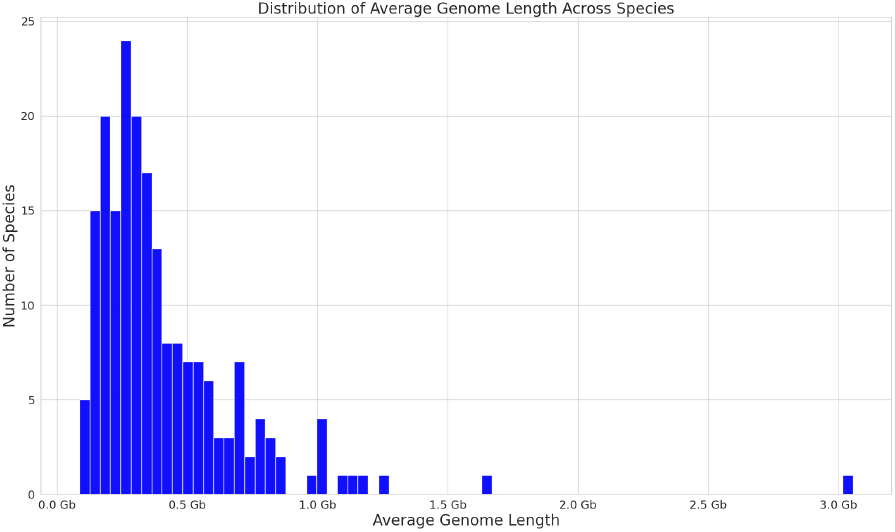
Distribution of genome sizes

**Figure 6:**
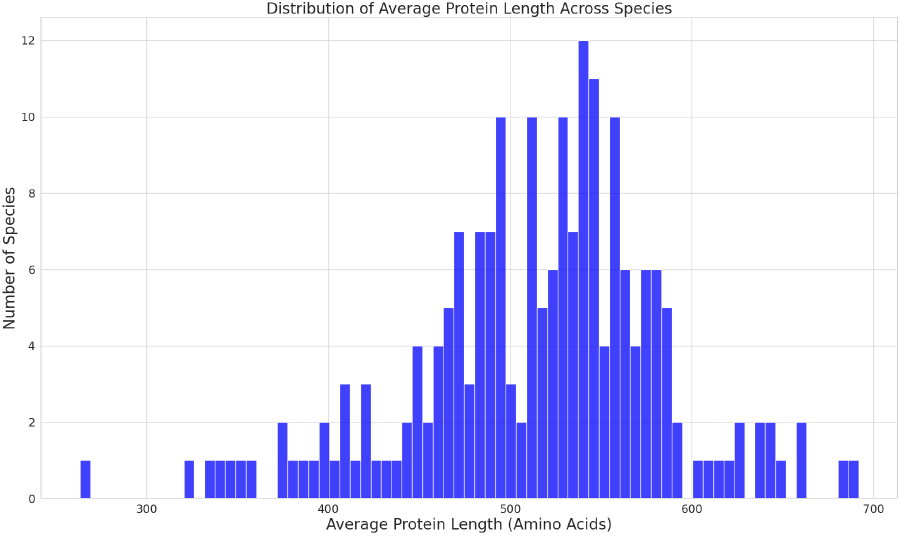
Distribution of predicted protein sizes

The 200 selected species are distributed across major insect clades as follows: **Endopterygota** (167 species), **Paraneoptera** (27), **Polyneoptera** (3), and **Palaeoptera** (3).

## Data Availability

We have deposited the dataset, including annotations, protein sequences, and OrthoFinder2 analysis in the figshare repositoryAll species-specific data are stored in separate `.tar.gz` archives. Each such archive contains four files: (i) predicted protein sequences including all isoforms, (ii) predicted protein sequences containing only the longest isoform per gene, (iii) a structural annotation file, and (iv) a structural annotation file decorated with Gene Ontology terms in `.gff3.gz` format. For example, `Drosophila_melanogaster.tar.gz` contains the four files `Drosophila_melanogaster{.faa,_longest.faa,.gtf,.gff3}`.

Additionally, for user convenience, the structural annotation files decorated with Gene Ontology terms (`.gff3.gz`) have also been uploaded separately to allow direct automated access. This results in duplication, as the `.gff3` files are available both individually and within the respective species archives. Furthermore, the results of OrthoFinder2 are provided in the archive `OrthoFinder2_Results.tar.gz`.

The genome sequences used as input for these annotations were obtained from NCBI. Details of the corresponding accession numbers, the specific BRAKER version used and whether the genomes had prior annotations are listed in Supplementary Table S1.

Regrettably, it was not possible to make the annotations produced in this study available via a third party annotation (TPA) submission to GenBank. The International Nucleotide Sequence Database Collaboration had announced in September 2024 [36] that third party annotations are not accepted anymore from January 2025.

## Technical Validation

### Gene Structure Accuracy

BRAKER3 was recently evaluated and compared against other pipelines in another study by comparing the predictions against reference annotations of 11 species [4]. In doing so, input protein evidence data from species closely related to the respective evaluation species were withheld. This was done to benchmark the application use case in which a new genome should be annotated. While Gabriel *et al*. [4] manually selected complete RNA-Seq libraries, the BRAKER-VARUS workflow used here automatically *samples* from RNA-Seq data with VARUS [11]. We therefore performed comparative tests of our results against those from the original BRAKER3 publication. We selected nine model organisms as benchmarks: *Danio rerio, Drosophila melanogaster, Gallus gallus, Mus musculus, Populus trichocarpa, Medicago truncatula, Parasteatoda tepidariorum, Arabidopsis thaliana*, and *Caenorhabditis elegans*. In addition, we evaluated variations in batch size and the number of RNA-Seq datasets. Unlike the original BRAKER3 approach, in which SRA datasets were manually selected prior to testing, the RNA data provision process in this study was fully automated. Below test aimed to demonstrate that the automatic selection of RNA-Seq datasets and their alignment to a reference genome does not compromise the quality of the results. The evaluated metrics were sensitivity, specificity, and the F1-score.

The results of the benchmarking are available in Table 1. Averaging over the nine species, the accuracy of the BRAKER-VARUS workflow with automatic RNA-Seq sampling from among all libraries in SRA is very close to the accuracy of BRAKER3 when complete libraries were chosen manually. Gene- and transcript-level accuracy are slightly better in the automated workflow, and exon-level accuracy is slightly worse, all differences are at most 0.6 percent points. We therefore conclude that our fully automated BRAKER-VARUS workflow, which only requires the binomial names as manually prepared input, is on average as good as a semi-automatic approach, where RNA-Seq libraries are prepared manually.

**Table 1:**
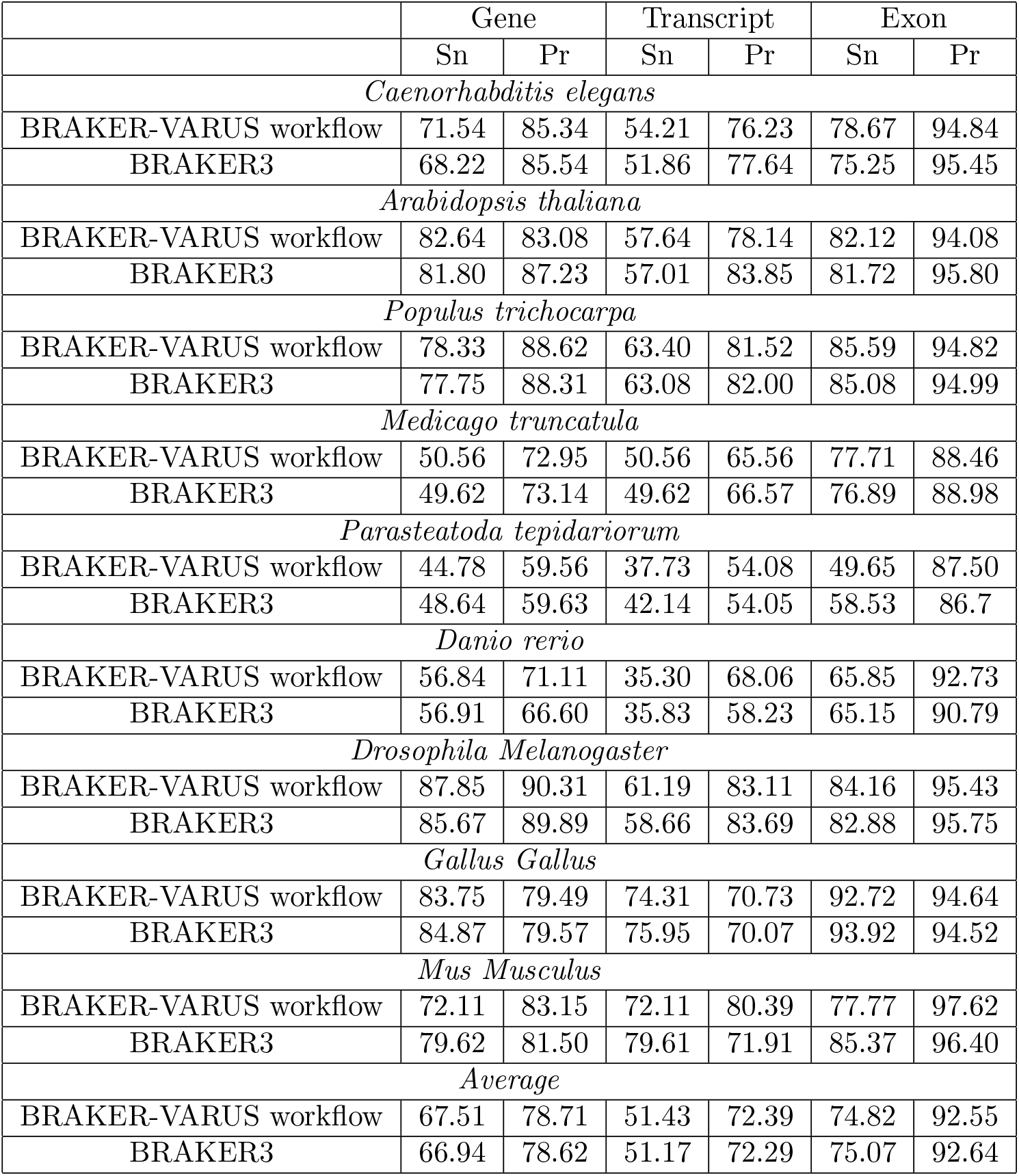
Gene Structure Accuracy (Sn=sensitivity, Pr=precision).

The following formulas were used to calculate the evaluation metrics:

- Sensitivity 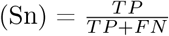, where *TP* is the number of true positives, and *FN* is the number of false negatives.
- Precision 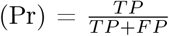, where *TP* is the number of true positives, and *FP* is the number of false positives.
- 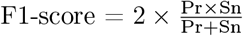.

### Proteome Completeness

To obtain one quality measure about proteome completeness, we can BUSCO [9]. For each species the predicted proteomes were evaluated using a lineage-specific dataset, such as hymenoptera_odb10, lepidoptera_odb10, diptera_odb10. For the selected genomes, BUSCO showed an annotation completeness level of at least 85%. The BUSCO results are shown in Supplementary Figure S1.

### Overprediction or Contamination

To obtain a second opinion on proteome completeness and to get information on overprediction or contamination, we run OMArk. OMArk reports an average completeness level of 92% and shows low contamination level (Supplementary Figure S2).

## Code Availability

The complete genome annotation workflow, including scripts and configurations for data processing, annotation, phylogenetic analysis with OrthoFinder, and Gene Ontology terms predictions with FANTASIA is available on GitHub at https://github.com/Gaius-Augustus/varus-braker/. Further details on the workflow and additional custom scripts can be found in the repository documentation.

## Acknowledgements

The work was funded in part by the Deutsche Forschungsgemeinschaft (DFG, German Research Foundation) – STA 1009/15-1. The computational analyses were performed using the HPC resources provided by Greifswald University and Münster University; for the latter, we thank the group of Erich Bornberg-Bauer. We also thank Elias Dohmen for providing the instructions to the HPC resources and discussing the OrthoFinder2 workflow for orthogroup analysis, and Gregor Bucher and Lena Reim for phylogeny.

## Author contributions statement

S.S. implemented and ran the workflow and analyzed the data. K.H. contributed to the creation of the workflow and structure annotation postprocessing. S.S. and M.S. designed the experiments. S.S. and M.S. wrote the draft manuscript. All authors reviewed, revised, and approved the manuscript.

