## Supplementary Material for "Annotation of 200 Insect Genomes with BRAKER for Consistent Comparisons across Species"

### Supplementary Materials

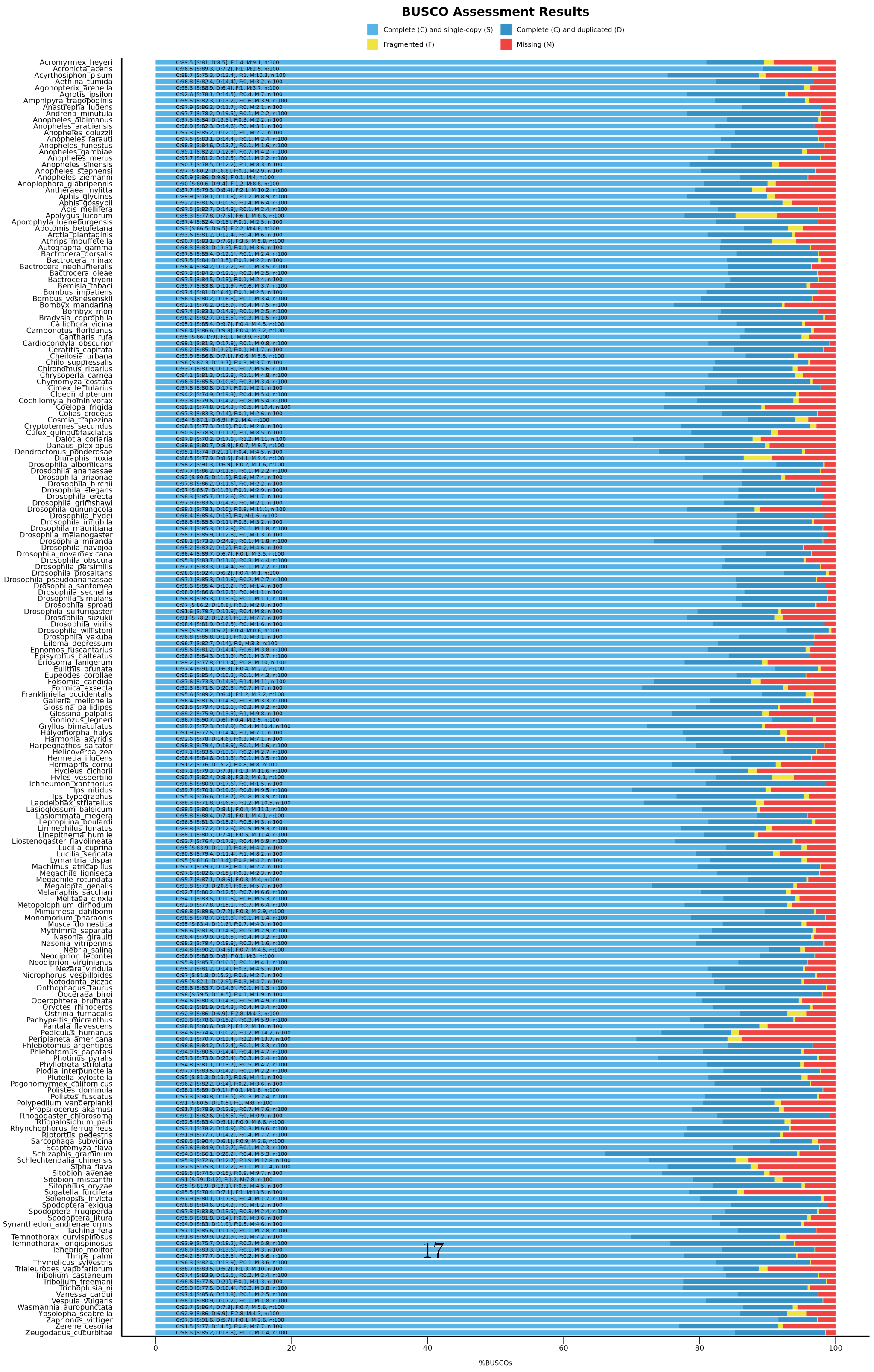


Supplementary Figure S1: BUSCO scoring table

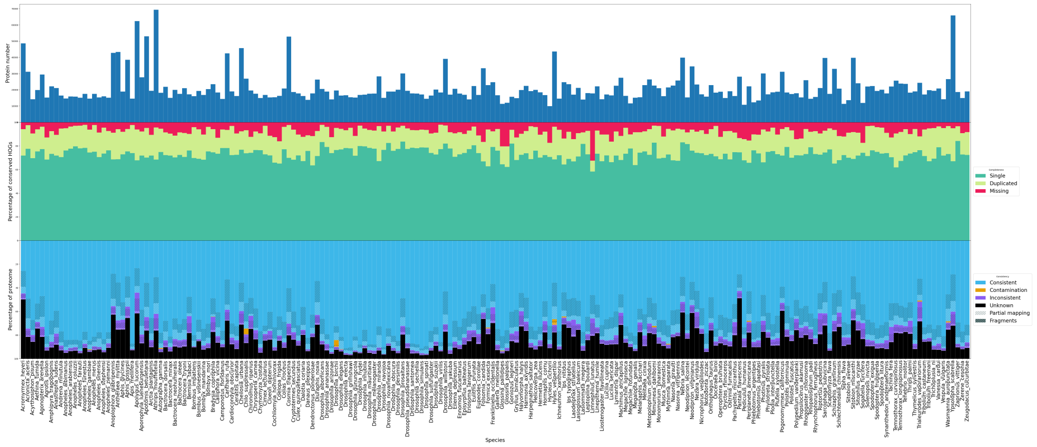


Supplementary Figure S2: OMArk scoring table

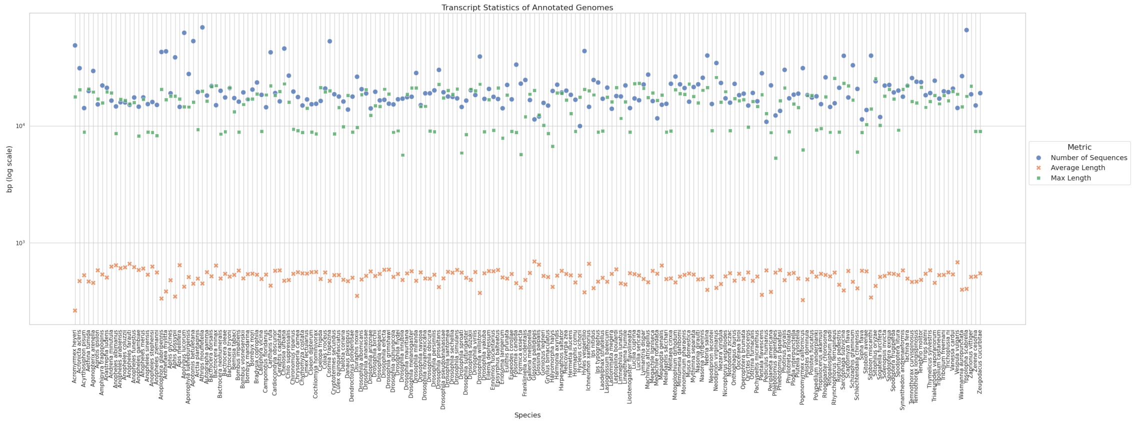


Supplementary Figure S3: Distribution of protein lengths and numbers

| **Species** | **Accession ID** | **BRAKER Version** | **Annotation Status (GenBank)** |
|---|---|---|---|
| *Acromyrmex heyeri* | GCA_017607565.1 | v2 | Annotated |
| *Acronicta aceris* | GCA_910591435.1 | v2 | Not Annotated |
| *Acyrthosiphon pisum* | GCF_005508785.2 | v3 | Annotated |
| *Aethina tumida* | GCF_024364675.1 | v3 | Annotated |
| *Agonopterix arenella* | GCA_927399405.1 | v2 | Not Annotated |
| *Agrotis ipsilon* | GCA_028554685.1 | v3 | Not Annotated |
| *Amphipyra tragopoginis* | GCA_905220435.1 | v3 | Not Annotated |
| *Anastrepha ludens* | GCF_028408465.1 | v3 | Not Annotated |
| *Andrena minutula* | GCA_929113495.1 | v3 | Not Annotated |

| | | | |
|---|---|---|---|
| *Anopheles albimanus* | GCF_013758885.1 | v3 | Annotated |
| *Anopheles arabiensis* | GCF_016920715.1 | v3 | Annotated |
| *Anopheles coluzzii* | GCF_943734685.1 | v3 | Annotated |
| *Anopheles farauti* | GCA_000473445.2 | v3 | Not Annotated |
| *Anopheles funestus* | GCF_943734845.2 | v3 | Annotated |
| *Anopheles gambiae* | GCF_000005575.2 | v3 | Not Annotated |
| *Anopheles merus* | GCF_017562075.2 | v3 | Annotated |
| *Anopheles sinensis* | GCA_000441895.2 | v3 | Annotated |
| *Anopheles stephensi* | GCF_013141755.1 | v3 | Annotated |
| *Anopheles ziemanni* | GCA_943734765.2 | v3 | Not Annotated |
| *Anoplophora glabripennis* | GCF_000390285.2 | v3 | Annotated |
| *Antheraea mylitta* | GCA_014332785.1 | v2 | Not Annotated |
| *Aphis glycines* | GCA_009761285.1 | v3 | Annotated |
| *Aphis gossypii* | GCF_020184175.1 | v2 | Annotated |
| *Apis mellifera* | GCF_003254395.2 | v3 | Annotated |
| *Apolygus lucorum* | GCA_009739505.2 | v2 | Annotated |
| *Aporophyla lueneburgensis* | GCA_932294355.1 | v3 | Not Annotated |
| *Apotomis betuletana* | GCA_932273695.1 | v2 | Not Annotated |
| *Arctia plantaginis* | GCA_902825455.1 | v3 | Annotated |
| *Athrips mouffetella* | GCA_947532105.1 | v2 | Not Annotated |
| *Autographa gamma* | GCA_905146925.1 | v3 | Not Annotated |
| *Bactrocera dorsalis* | GCF_023373825.1 | v3 | Annotated |
| *Bactrocera minax* | GCA_021498325.1 | v3 | Not Annotated |
| *Bactrocera neohumeralis* | GCF_024586455.1 | v3 | Annotated |
| *Bactrocera oleae* | GCF_001188975.3 | v3 | Annotated |
| *Bactrocera tryoni* | GCF_016617805.1 | v3 | Annotated |
| *Bemisia tabaci* | GCF_001854935.1 | v3 | Annotated |
| *Bombus impatiens* | GCF_000188095.3 | v3 | Annotated |
| *Bombus vosnesenskii* | GCF_011952255.1 | v3 | Annotated |
| *Bombyx mandarina* | GCF_003987935.1 | v3 | Annotated |
| *Bombyx mori* | GCF_014905235.1 | v3 | Annotated |
| *Bradysia coprophila* | GCA_014529535.2 | v3 | Annotated |
| *Calliphora vicina* | GCA_958450345.1 | v3 | Not Annotated |
| *Camponotus floridanus* | GCF_003227725.1 | v3 | Annotated |
| *Cantharis rufa* | GCA_947369205.1 | v2 | Not Annotated |
| *Cardiocondyla obscurior* | GCA_019399895.1 | v3 | Not Annotated |
| *Ceratitis capitata* | GCF_000347755.3 | v3 | Annotated |
| *Cheilosia urbana* | GCA_946477585.1 | v2 | Not Annotated |
| *Chilo suppressalis* | GCA_902850365.2 | v3 | Annotated |
| *Chironomus riparius* | GCA_917627325.4 | v3 | Annotated |
| *Chrysoperla carnea* | GCF_905475395.1 | v3 | Annotated |
| *Chymomyza costata* | GCA_018150985.1 | v3 | Not Annotated |
| *Cimex lectularius* | GCF_000648675.2 | v3 | Annotated |
| *Cloeon dipterum* | GCA_949628265.1 | v3 | Annotated |

| | | | |
|---|---|---|---|
| *Cochliomyia hominivorax* | GCA_004302925.2 | v3 | Not Annotated |
| *Coelopa frigida* | GCA_017309665.1 | v3 | Not Annotated |
| *Colias croceus* | GCF_905220415.1 | v3 | Annotated |
| *Cosmia trapezina* | GCA_905163495.3 | v2 | Not Annotated |
| *Cryptotermes secundus* | GCF_002891405.2 | v3 | Annotated |
| *Culex quinquefasciatus* | GCF_015732765.1 | v3 | Annotated |
| *Dalotia coriaria* | GCA_025399875.2 | v3 | Not Annotated |
| *Danaus plexippus* | GCF_009731565.1 | v3 | Not Annotated |
| *Dendroctonus ponderosae* | GCF_020466585.1 | v3 | Annotated |
| *Diuraphis noxia* | GCF_001186385.1 | v3 | Annotated |
| *Drosophila albomicans* | GCF_009650485.2 | v2 | Annotated |
| *Drosophila ananassae* | GCF_017639315.1 | v3 | Annotated |
| *Drosophila arizonae* | GCF_001654025.1 | v3 | Annotated |
| *Drosophila birchii* | GCA_008042755.1 | v3 | Annotated |
| *Drosophila elegans* | GCF_018152505.1 | v3 | Annotated |
| *Drosophila erecta* | GCF_003286155.1 | v3 | Annotated |
| *Drosophila grimshawi* | GCF_018153295.1 | v3 | Annotated |
| *Drosophila gunungcola* | GCF_025200985.1 | v3 | Annotated |
| *Drosophila hydei* | GCF_003285905.1 | v3 | Annotated |
| *Drosophila innubila* | GCA_004354385.2 | v3 | Annotated |
| *Drosophila mauritiana* | GCF_004382145.1 | v3 | Annotated |
| *Drosophila melanogaster* | GCF_000001215.4 | v3 | Annotated |
| *Drosophila miranda* | GCF_003369915.1 | v3 | Annotated |
| *Drosophila navojoa* | GCF_001654015.2 | v3 | Annotated |
| *Drosophila novamexicana* | GCF_003285875.2 | v2 | Annotated |
| *Drosophila obscura* | GCF_018151105.1 | v3 | Annotated |
| *Drosophila persimilis* | GCF_003286085.1 | v3 | Annotated |
| *Drosophila prosaltans* | GCA_018151275.1 | v2 | Not Annotated |
| *Drosophila pseudoananassae* | GCA_018153035.1 | v3 | Annotated |
| *Drosophila santomea* | GCF_016746245.2 | v3 | Annotated |
| *Drosophila sechellia* | GCF_004382195.2 | v3 | Annotated |
| *Drosophila simulans* | GCF_016746395.2 | v3 | Annotated |
| *Drosophila sproati* | GCA_018904355.1 | v3 | Not Annotated |
| *Drosophila sulfurigaster* | GCA_023558435.1 | v3 | Annotated |
| *Drosophila suzukii* | GCF_013340165.1 | v3 | Annotated |
| *Drosophila virilis* | GCF_003285735.1 | v3 | Annotated |
| *Drosophila willistoni* | GCF_018902025.1 | v2 | Annotated |
| *Drosophila yakuba* | GCF_016746365.2 | v3 | Annotated |
| *Eilema depressum* | GCA_914767945.1 | v3 | Not Annotated |
| *Ennomos fuscantarius* | GCA_905220475.3 | v3 | Not Annotated |
| *Episyrphus balteatus* | GCF_945859705.1 | v3 | Not Annotated |
| *Eriosoma lanigerum* | GCA_013282895.1 | v3 | Not Annotated |
| *Eulithis prunata* | GCA_918843925.1 | v2 | Not Annotated |
| *Eupeodes corollae* | GCF_945859685.1 | v3 | Not Annotated |

| | | | |
|---|---|---|---|
| *Folsomia candida* | GCF_002217175.1 | v3 | Annotated |
| *Formica exsecta* | GCF_003651465.1 | v3 | Annotated |
| *Frankliniella occidentalis* | GCF_000697945.3 | v3 | Annotated |
| *Galleria mellonella* | GCF_026898425.1 | v3 | Annotated |
| *Glossina pallidipes* | GCA_000688715.1 | v3 | Not Annotated |
| *Glossina palpalis* | GCA_000818775.1 | v3 | Not Annotated |
| *Goniozus legneri* | GCA_003055095.1 | v2 | Not Annotated |
| *Gryllus bimaculatus* | GCA_017312745.1 | v3 | Not Annotated |
| *Halyomorpha halys* | GCA_000696795.3 | v3 | Not Annotated |
| *Harmonia axyridis* | GCF_914767665.1 | v3 | Annotated |
| *Harpegnathos saltator* | GCF_003227715.2 | v3 | Annotated |
| *Helicoverpa zea* | GCF_022581195.2 | v3 | Annotated |
| *Hermetia illucens* | GCF_905115235.1 | v3 | Annotated |
| *Hormaphis cornu* | GCA_017140985.1 | v3 | Not Annotated |
| *Hycleus cichorii* | GCA_013841215.1 | v3 | Not Annotated |
| *Hyles vespertilio* | GCA_009982885.2 | v2 | Not Annotated |
| *Ichneumon xanthorius* | GCA_917499995.1 | v3 | Not Annotated |
| *Ips nitidus* | GCA_018691245.2 | v3 | Not Annotated |
| *Ips typographus* | GCA_016097725.1 | v3 | Not Annotated |
| *Laodelphax striatellus* | GCA_017141395.1 | v3 | Annotated |
| *Lasioglossum baleicum* | GCA_022376115.1 | v2 | Not Annotated |
| *Lasiommata megera* | GCA_928268935.1 | v3 | Not Annotated |
| *Leptopilina boulardi* | GCF_019393585.1 | v3 | Not Annotated |
| *Limnephilus lunatus* | GCA_917563855.2 | v3 | Not Annotated |
| *Linepithema humile* | GCF_000217595.1 | v2 | Annotated |
| *Liostenogaster flavolineata* | GCA_025433975.1 | v3 | Not Annotated |
| *Lucilia cuprina* | GCF_022045245.1 | v3 | Annotated |
| *Lucilia sericata* | GCF_015586225.1 | v3 | Annotated |
| *Lymantria dispar* | GCA_016802235.1 | v3 | Not Annotated |
| *Machimus atricapillus* | GCA_933228815.1 | v3 | Not Annotated |
| *Megachile ligniseca* | GCA_945859555.1 | v3 | Not Annotated |
| *Megachile rotundata* | GCF_000220905.1 | v3 | Annotated |
| *Megalopta genalis* | GCF_011865705.1 | v3 | Annotated |
| *Melanaphis sacchari* | GCF_002803265.2 | v3 | Annotated |
| *Melitaea cinxia* | GCF_905220565.1 | v3 | Annotated |
| *Metopolophium dirhodum* | GCA_019925205.1 | v3 | Not Annotated |
| *Mimumesa dahlbomi* | GCA_917499265.3 | v2 | Not Annotated |
| *Monomorium pharaonis* | GCF_013373865.1 | v3 | Annotated |
| *Musca domestica* | GCF_000371365.1 | v3 | Annotated |
| *Mythimna separata* | GCA_029852925.1 | v3 | Not Annotated |
| *Nasonia giraulti* | GCA_016647725.1 | v3 | Not Annotated |
| *Nasonia vitripennis* | GCF_009193385.2 | v3 | Annotated |
| *Nebria salina* | GCA_944039245.1 | v2 | Not Annotated |
| *Neodiprion lecontei* | GCF_021901455.1 | v3 | Annotated |

| | | | |
|---|---|---|---|
| *Neodiprion virginianus* | GCF_021901495.1 | v2 | Annotated |
| *Nezara viridula* | GCA_928085145.1 | v3 | Annotated |
| *Nicrophorus vespilloides* | GCF_001412225.1 | v3 | Annotated |
| *Notodonta ziczac* | GCA_918843915.1 | v3 | Not Annotated |
| *Onthophagus taurus* | GCF_000648695.1 | v3 | Annotated |
| *Ooceraea biroi* | GCF_003672135.1 | v3 | Annotated |
| *Operophtera brumata* | GCA_932527175.1 | v3 | Annotated |
| *Oryctes rhinoceros* | GCA_020654165.1 | v3 | Not Annotated |
| *Ostrinia furnacalis* | GCF_004193835.2 | v3 | Not Annotated |
| *Pachypeltis micranthus* | GCA_020466155.1 | v3 | Not Annotated |
| *Pantala flavescens* | GCA_020796165.1 | v2 | Not Annotated |
| *Pediculus humanus* | GCF_000006295.1 | v3 | Not Annotated |
| *Periplaneta americana* | GCA_025594305.2 | v3 | Annotated |
| *Phlebotomus argentipes* | GCA_947086385.1 | v3 | Not Annotated |
| *Phlebotomus papatasi* | GCF_024763615.1 | v3 | Not Annotated |
| *Photinus pyralis* | GCF_008802855.1 | v3 | Annotated |
| *Phyllotreta striolata* | GCA_918026865.1 | v3 | Annotated |
| *Plodia interpunctella* | GCF_027563975.1 | v3 | Annotated |
| *Plutella xylostella* | GCF_932276165.1 | v3 | Annotated |
| *Pogonomyrmex californicus* | GCA_024349325.1 | v3 | Not Annotated |
| *Polistes dominula* | GCF_001465965.1 | v2 | Annotated |
| *Polistes fuscatus* | GCF_010416935.1 | v3 | Annotated |
| *Polypedilum vanderplanki* | GCA_018290095.1 | v3 | Annotated |
| *Propsilocerus akamusi* | GCA_018397935.1 | v3 | Not Annotated |
| *Rhogogaster chlorosoma* | GCA_944452935.1 | v3 | Not Annotated |
| *Rhopalosiphum padi* | GCA_020882245.1 | v3 | Not Annotated |
| *Rhynchophorus ferrugineus* | GCA_030347505.1 | v3 | Annotated |
| *Riptortus pedestris* | GCA_019009955.1 | v3 | Not Annotated |
| *Sarcophaga subvicina* | GCA_936449025.2 | v2 | Not Annotated |
| *Scaptomyza flava* | GCA_030179655.1 | v3 | Not Annotated |
| *Schizaphis graminum* | GCA_020882235.1 | v3 | Not Annotated |
| *Schlechtendalia chinensis* | GCA_019022885.1 | v3 | Not Annotated |
| *Sipha flava* | GCF_003268045.1 | v3 | Annotated |
| *Sitobion avenae* | GCA_019425605.1 | v3 | Not Annotated |
| *Sitobion miscanthi* | GCA_008086715.1 | v2 | Not Annotated |
| *Sitophilus oryzae* | GCF_002938485.1 | v3 | Annotated |
| *Sogatella furcifera* | GCA_017141385.1 | v3 | Not Annotated |
| *Solenopsis invicta* | GCF_016802725.1 | v3 | Annotated |
| *Spodoptera exigua* | GCA_902829305.4 | v3 | Annotated |
| *Spodoptera frugiperda* | GCF_023101765.2 | v3 | Annotated |
| *Spodoptera litura* | GCF_002706865.2 | v3 | Annotated |
| *Synanthedon andrenaeformis* | GCA_936446665.2 | v3 | Not Annotated |
| *Tachina fera* | GCA_905220375.1 | v3 | Not Annotated |
| *Temnothorax curvispinosus* | GCF_003070985.1 | v3 | Annotated |

| | | | |
|---|---|---|---|
| *Temnothorax longispinosus* | GCA_030848805.1 | v3 | Annotated |
| *Tenebrio molitor* | GCA_907166875.3 | v3 | Annotated |
| *Thrips palmi* | GCF_012932325.1 | v3 | Annotated |
| *Thymelicus sylvestris* | GCA_911387775.1 | v3 | Not Annotated |
| *Trialeurodes vaporariorum* | GCA_011764245.1 | v2 | Not Annotated |
| *Tribolium castaneum* | GCF_000002335.3 | v3 | Annotated |
| *Tribolium freemani* | GCA_939628115.1 | v3 | Not Annotated |
| *Trichoplusia ni* | GCF_003590095.1 | v3 | Annotated |
| *Vanessa cardui* | GCA_905220365.2 | v3 | Annotated |
| *Vespula vulgaris* | GCF_905475345.1 | v3 | Annotated |
| *Wasmannia auropunctata* | GCF_000956235.1 | v2 | Annotated |
| *Ypsolopha scabrella* | GCA_910592155.1 | v2 | Not Annotated |
| *Zaprionus vittiger* | GCA_018904025.1 | v2 | Not Annotated |
| *Zerene cesonia* | GCF_012273895.1 | v3 | Annotated |
| *Zeugodacus cucurbitae* | GCF_028554725.1 | v3 | Not Annotated |

Supplementary Table S1: Species accession IDs and annotation status
